# A telomere-to-telomere (T2T) pig genome assembly reveals Y chromosome diversity and structural variations of Wuzhishan pigs

**DOI:** 10.64898/2026.04.23.720499

**Authors:** Yuwei Ren, Feng Wang, Xinjian Li, Guangliang Liu, Ruiping Sun, Xinli Zheng, Yan Zhang, Ruiyi Lin, Xuyang Lu, Linlin Chen, Wenshui Xin, Yanli Fei, Zhe Chao

## Abstract

**Backgroud:** Wuzhishan (WZS) pigs are native to Hainan Province of China, and serve as both important agricultural resources and biomedical models. Although the published WZS pig genome (T2T-pig1.0) even achieving telomere-to telomere (T2T) completeness, substantial genetic diversity still exists within the same pig breed, another WZS pig genome named WZS-T2T was assembled in this study.

**Results:** Multiple sequencing data were used to assemble genome, and finally yielded a ∼2.68 Gb telomere-to-telomere genome, with N50 length ∼142.87 Mb, and annotated protein coding genes of 23,100. Compared to T2T-pig1.0, QV and BUSCO value was higher, and the Y chromosome (ChrY) length was longer in WZS-T2T than that of T2T-pig1.0. ChrY of two WZS pigs shared 11 genes, including sex differentiation-related genes of *SHOX*, *PRKX*, and *DDX3X*, and *SRY*; however, energy metabolism gene *SLC25A4* and the macrophage-related receptor gene *CSF2RA* of ChrY were specific to WZS-T2T. An inversion SV on chromosome 10 with length ∼33.86 Mb was identified between two WZS pigs, and three proofs were proposed for proving the accuracy sequence orientation of WZS-T2T.The genetic diversity was consistent with LD decay speed in population different analysis. WZS pigs exhibited higher genetic diversity than other four pig populations (Tunchang pigs, Yuxi black pigs, Large White pig, and Duroc pigs) examined in this study, and presented slower LD decay compared to other four breeds.

**Conclusions:** Therefore, WZS-T2T provided a higher-quality assembly, and potential advantages of both agricultural production and biomedical targets for WZS pigs.

## Introduction

Pigs are vital to human society, supplying high-quality protein, and serving as models for biomedical research [1, 2]. Chinese indigenous pig breeds have evolved diverse adaptive traits in response to local environments. Remarkably, pigs increasingly favored as animal models in the developing of treatments and vaccines due to their physiological, immunological, and genetic similarities to humans [3]. For example, WZS pigs with streptozotocin induced diabetes disease could accurately mimic the pathological symptoms of human diabetes, and that pancreatic islet beta-cell regeneration occurred in an adult WZS pig [4]. WZS pigs were engineered with deletion of the gene encoding glycoprotein alpha-galactosyltransferase 1 (*GGTA1*) with the goal of improving their utility as a source of biomedical tissues [5]. Heart valve, vascular grafts [6] and coronary stent were developed for preclinical trials [7]. In 2022, the first pig-to-human heart transplant was performed at the University of Maryland, extending survival to ∼60 days [8, 9]. More recent cases highlight the rapid progress of xenotransplantation and its potential to extend human life [10].

Wuzhishan (WZS) pigs, a minipig native to Hainan Province of China, are characterized by flavorful meat, distinctive coat colors of black back with a white belly, and an inverted triangle on the forehead. The pigs are value as biomedical models, particularly in human disease research [4]. Recently, numerous genome assemblies for pigs have been published [11–13], and WZS pig genome (GCA_048338725.1) named T2T-pig1.0 even achieving telomere-to telomere (T2T) completeness [14]. However, as considerable genetic diversity still exists within cultivated breed, we assembled a WZS pig genome (GWHHNIC00000000.1) named WZS-T2T in this study to provide a higher-quality genome of WZS pigs by comparing genetic characteristics of two WZS pig genomes, including the differences of genome assembly and quality, annotations, Y chromosome, genome structural variations (SVs).

## Methods

### Sample collection and genome sequencing

One 8-month-old male Wuzhishan pig was obtained from the Wuzhishan Preservation Factory (Chengmai city, Hainan Province). The pig was farmed with ad libitum food and water for one week before slaughter. After 24 hours of feed withdrawal, it was transferred to the slaughterhouse in the HAAS Yongfa pig experimental facility for slaughter. The pig was stunned by electric shock and exsanguinated, flushed, and split without regaining consciousness. Blood, heart, liver, spleen, kidney, and longissimus dorsi muscle (LDM) samples were collected.

All the tissues were snap-frozen in liquid nitrogen and stored on dry ice, and then sent to Novogene Company (Beijing, China) and Huaming Company (Wuhan, China) for sequencing. Genomic DNA was extracted from LDM and was used to construct PCR-free 350-bp insert libraries, and performed next generation sequencing (NGS) on Illumina NOVA6000 platform. Blood-derived DNA was used for High-Fidelity Sequencing (HiFi) and Oxford Nanopore Technologies (ONT) libraries. HiFi reads were generated on PacBio Sequel IIe system in circular consensus sequencing (ccs) mode, while ultra-long reads were obtained via ONT sequencing. High-throughput/resolution chromosome conformation capture (Hi-C) library was prepared by in situ crosslinking nuclear DNA, digestion with restriction enzymes, end biotinylation, dilution, and random ligation. The ligated DNA fragments were enriched, polished, and sequenced. RNA was extracted from the pooled heart, liver, spleen, kidney, and LDM samples and sequenced on the Illumina NOVA6000.

The Illumina raw data was filtered with low quality reads and adapters using fastp (version 0.21.0) [15]. K-mer analysis was further performed with Jellyfish (version 2.3.0) to estimate genome size, heterozygosity, and repeat content. PacBio raw data of were processed with ccs (version 6.0.0) under the parameters “–min-passes 3 –min-snr 2.5 –top-passes 60”. ONT raw data were transferred to fastq file though base calling by Guppy software (version 1.2.0) [16], and reads≥Q7 were filtered using Filtlong (version 0.2.4) to obtain “pass” reads with sequence length≥10Kb, and N50 ∼81 Kb. Hi-C data were trimmed low quality reads and adapters with fastp (version 0.21.0), and removed unmapped reads, self-circle reads, and dangling end reads with HICUP (version 0.8.0) [17].

High-quality HiFi reads, filtered ONT ultra-long reads (N50 ∼81 Kb) and Hi-C data were integrated for assembly using Hifiasm (version 0.24.0-r702) [18]. The initially high-quality assembly combined HiFi, ONT and Hi-C data. Primary contigs were purged of redundancy and heterozygosity by Purge_dups with default parameters (version 1.2.5) [19], and anchored to Hi-C data for phasing and scaffolding by ALLHiC (version 0.9.13) [20], and processed with Juicer (version 1.6) and 3D-DNA (version 180419), with manual refinement in Juicebox (version 1.11.08) [21]. Gaps were initially filled as 100 Ns, and unanchored contigs were designated as chrUnn. Gaps were further resolved using ONT reads with TGS-GapCloser (version 1.2.1) [22], and polished with Racon [23], yielding a telomere-to-telomere (T2T) WZS pig genome assembly.

The genome quality was tested by mapping Illumina reads, HiFi data, ONT data back to the genome sequences using BLASTN (version 2.3.0+) [24], and calculated QV with Merqury (version 1.3) and GCI of HiFi and ONT data with GCI (Genome Continuity Inspector, version 1.0) [25], and BUSCO values based on BUSCO (Benchmarking Universal Single-Copy Orthologs, version 5.3.2).

### Identification of telomere and centromere

Telomere detection was performed by mapping ONT reads to the assembled genome using Winnowmap (version 1.11) [26]. Reads containing the tandem repeat sequence ‘CCCTAA’/‘TTAGGG’ with the highest repeat counts were defined as telomere references, while other matched reads were designated as queries. Telomere repeats were quantified using the TeloExplorer module within quarTeT (version 1.2.0) [27].

Telomeres were reassembled from reference and query reads using Medaka (version 2.0.1) (https://github.com/nanoporetech/medaka). The reconstructed telomeres were then mapped to draft genome, and tandem repeats with≤80% identity at chromosomes ends were replaced using MUMmer (version 3.23) [28].

Centromere candidates were predicted using the CentroMiner module within quarTeT (version 1.2.0) [27]. TRF (version 4.09.1) was used to screen tandem repeats (TRs) across chromosomes, and representative monomers were selected based on period and copy number. Redundant monomers were removed by clustering, and the remaining centromere monomers were aligned to chromosomes with cd-hit (version 4.8.1) (https://github.com/weizhongli/cdhit). Candidate centromeres were further refined based on sequence identity and mapping length. Centromere distribution and TRs density were analyzed and visualized using R package RIdeogram (version 0.2.2) [29].

### Annotation of the Wuzhishan pig genome

Repeat sequences included transposable elements (TEs) and tandem repeats (TRs), which were annotated through both *de novo* and homology-based approaches. A *de novo* TE library was generated using RepeatModeler (version 2.0.5) [30]. Long terminal repeats (LTRs) were identified with LTR_FINDER (version 1.07) and LTRharvest (version 1.62) [31], followed by redundancy filtering using HiTE (version 3.3.3) [32] to obtain non-redundant LTRs. Terminal inverted repeats (TIRs) were classified by GenericRepeatFinder (version 1.0.2) [33] and TIR-Learner (version 3.0.7) [34]. Helitron was predicted using Heliano (version 1.3.1) and HelitronScanner (version 1.0). These sequences were combined with the *de novo* TE library, resulting in a high-quality TE library. Both the *de novo* TE library, and the RepBase database (version 20181026) [35] and Dfam database (version 20250505) were combined, and filtered low-confidence level sequences. Unknown TEs were classified by TEclass (version 2.1.3) [36] and NeuralTE (version 1.0.1) [37]. All TE predictions were integrated, and redundancy was removed to yield the final TE set, and annotated using RepeatProteinMask within RepeatMasker (version 4.2.0). TRs were predicted with TRF (version 4.10.0rc2) and MISA (version 2.1), and simple sequence repeats (SSRs) were detected using MIcroSAtellite identification tool (MISA) (http://pgrc.ipk-gatersleben.de/misa/) [38].

Gene annotation incorporated transcriptome, *de novo* and homology-based predictions. Transcriptome-based annotation was performed using Illumina RNA-seq from pooled samples (heart, liver, spleen, kidney, longissimus dorsi muscle (LDM). Reads were mapped with HISAT2 (version 2.2.1). assembled with StringTie (version 2.2.1) [39–41], and open reading frame (ORF) were predicted with TransDecoder (version 5.7.1) [42]. *De novo* predictions were generated with BRAKER (version: 3.0.8) [43], ANNEVO (version: 2.2) and Helixer (version 0.3.6) [44] using the repeat-masked genome. For homology-based annotation, protein sequences from *Sus scrofa* 11.1 (GCF_000003025.6, Duroc), Jinhua pigs (https://alphaindex.zju.edu.cn/ALPHADB/download.html), Rongchang pigs (https://ngdc.cncb.ac.cn/gwh/Assembly/92155/show), and Min pigs (https://ngdc.cncb.ac.cn/gwh/Assembly/92156/show) were mapped to the WZS pig genome using Miniprot (version 0.18). Finally, results from all three approaches were integrated using MAKER (version 3.01.03) [45] and EVidenceModele (version 2.1.0).

Functional annotation was conducted by aligning predicted proteins to the NCBI NR database (Non-Redundant Protein Sequence Database) using Diamond (version 2.0.11.149) [46] and to UniProt using BLASTP (version 2.3.0+) [24], with thresholds of *E*-value ≤ 10^-5^, coverage ≥ 80% and identity ≥ 90%. Additional annotations were obtained from InterPro (version 107.0) and Pfam databases (version 38.0) using InterProScan (version5.47-82.0) [47]. GO (Gene Ontology) and KEGG (Kyoto Encyclopedia of Genes and Genomes) [48] pathway annotations were assigned with eggNOG-mapper (version 2.1.10) [49], while orthologous group classification was conducted with the KOG databased using Diamond (version 2.0.11.149) [46]. Gene expression profiles from RNA-seq were also integrated. Genes were retained if supported by at least two independent sources of evidence.

### Evolutionary analysis

Orthologous and paralogous genes were identified across six mammalian species within two pig breeds, including the WZS pig genome (GWHHNIC00000000.1) named WZS-T2T, Homo sapiens (GCF_000001405.40), *Sus scrofa* 11.1 (Duroc, GCF_000003025.6), Bos taurus (GCF_002263795.3), Canis lupus (GCA_011100685.1), Equus caballus (GCA_041296265.1), Ovis aries (GCA_016772045.2), using OrthoFinder (version 2.5.5) with default parameters [50, 51]. Orthologous genes (orthologs) for each species were selected to construct phylogenetic tree. Protein sequences of orthologs were aligned with MUSCLE (version 5) [52]with default parameters to concatenated into one supergene sequences for each species, and used to generate a phylogenetic tree using RAxML (version 8.2.10) by the algorithm of maximum likelihood (ML) [53]. Divergence times were estimated by PAML’s mcmctree (version 4.10.7) [54], calibrated against speciation times from TimeTree (http://www.timetree.org/) with most recent common ancestor (TMRCA) [55]. The tree was visualized with MCMCTreeR (version 1.1) [56]. Gene family expansion and contraction were analyzed with CAFÉ (version 5) [57]. Genomes collinearity was analyzed using MCScanX (version 1.0.0) between the WZS-T2T and the previously published WZS pigs genome (GCA_048338725.1) named T2T-pig1.0 [58], and the six pig breeds (WZS-T2T, Jinhua pigs, and Chenghua pigs) and *Homo sapiens* with default parameters.

### Genome structural variations (SVs) analysis

Structural variants (SVs) were identified by aligning the WZS-T2T and T2T-pig1.0 using MUMmer (version 3.23). Collinearity blocks, structural rearrangement (inversions, translocations, duplications), insertions, deletions, highly diverged regions were detected by SyRI (version: 1.6). The insertions with the length≥50 bp were defined as presence, and the deletions with the length≥50 bp were defined as absence, while highly diverged regions and not aligned regions were defined as presence/absence variation (PAV). The annotation of the SVs was using ANNOVAR (version 2025Mar02) [59].

Using the minimap2 software (version 2.28-r1209) (Li 2021), the HiFi and ONT data were aligned to the genome. The resulting alignment files (BAM and PAF) were used to visualize read coverage across chromosomes, with a focus on the inversion breakpoint position on chromosome 10.

### Y Chromosome analysis

Collinearity between the X and Y chromosomes was assessed with MUMmer (version 3.23). Sequence types and counts within the pseudoautosomal regions (PARs) were analyzed, and SVs and single nucleotide polymorphisms (SNPs) were identified using SyRI (version 1.6) [60]). Male-specific regions on Y (MSY) chromosome were identified by their position and length with SyRI (version 1.6). Syntenic regions between two WZS pigs (WZS-T2T, T2T-pig1.0) were analyzed by minimap2 software (version 2.28-r1209) [61], and visualized using R package RIdeogram (version 0.2.2) [29].

### Whole genome sequencing analysis

#### Sample collection and datasets

Population difference analysis were performed from 205 individuals, including 147 samples sequenced in this study, and 58 samples downloaded from NCBI. Ear tissue samples were collected from 147 pigs across five populations: 30 Wuzhishan (WZS) pigs (4 months old), 30 Tunchang (TC) pigs (6 months old), 29 Yuxi black (YX) pigs (8 months old), 30 Large White (LW) pigs (2 months old), and 28 Duroc pigs (8 months old); and data of 32 LW pigs and 26 WZS pigs were downloaded from NCBI accession PRJNA994680. Ears samples of WZS, TC, LW, and Duroc pigs were obtained from Hainan Province, while YX pigs were collected from a farm in Henan Province. All ear tissues were preserved in 100% ethanol and stored at −80°C for downstream genomic analyses.

#### Resequencing and variant calling

Genomic DNA was extracted using a DNeasy Blood & Tissue Kit (Qiagen, Shanghai, China). DNA quality and concentration were verified by agarose gel electrophoresis and measured using a Qubit 4.0 Fluorometer (Thermo Fisher Scientific, 739256). Approximately 1 μg of DNA from each individual was used to prepare paired-end sequencing libraries with an average insert size of ∼350 bp. Libraries quality was checked using an Agilent 2100 Bioanalyzer (Agilent, Waldbronn, Germany) before sequencing on the Illumina Nova 6000 platform, following standard protocols at BioMarker Company (Beijing, China).

Raw sequencing reads were first processed with fastp (version 0.21.0) to remove low-quality bases and adapters, and then aligned to the WZS pig reference genome using Burrows-Wheeler-Aligner (BWA) (BWA-MEM, version 0.7.17) (https://sourceforge.net/projects/bio-bwa/files/). The resulting alignments were sorted with SAMtools (version 1.20) [62], and PCR duplicate reads were eliminated using the MarkDuplicates module in GATK (version 4.2.2.0) (https://gatk.broadinstitute.org/hc/en-us) [63]. SNPs and small insertions/deletions (indels) were called using the Haplotyper module in Sentieon software (version 202112.05). retaining only sites with sequencing depth greater than 6×. Combined gVCF files were processed with GenotypeGVCFs in Genome Analysis Toolkit (GATK) (version 4.2.2.0) to obtain population-level SNPs and indels. SNPs were extracted using SelectVariants with the parameter “-select-type-to-include SNP” and further filtered with VariantFiltering using the thresholds: QualByDepth (QD)<2.0, RMSMappingQuality (MQ)<40.0, FisherStrand (FS)>60.0, StrandOddsRatio (SOR)>3.0, MappingQualityRankSumTest (MQRankSum)<−12.5, and ReadPosRankSumTest (ReadPosRankSum)<−8.0 [64]. Functional annotation of retained SNPs was performed with SnpEff based on the WZS pig reference genome [65].

#### Population difference analysis

Genetic distance was assessed by computing an identity-by-state (IBS) matrix using PLINK (version 2.0) [66], and a phylogenetic tree was generated using the neighbor-joining (NJ) algorithm in Phylip (version 3.698) [67]. Principal component analysis (PCA) was performed using the R package FactoMineR (version 2.12) to identify population clustering patterns.

High-confidence SNPs were subsequently used to quantify genetic diversity and population differentiation. Nucleotide diversity (π) was calculated with VCFtools (version 0.1.17) [68] in 1 Mb non-overlapping windows. Population divergence was evaluated using Weir and Cockerham’s fixation index (*F_ST_*) [69] in 300 Kb windows via VCFtools (version 0.1.17). Linkage disequilibrium (LD) decay was evaluated using the squared correlation coefficient (r²) at a maximum distance of 300 kb with PopLDdecay (version 3.41) [70].

## Results

### Telomere-to-telomere genome assembly

We integrated multiple sequencing technologies, including Illumina, PacBio, ONT, and Hi-C to assemble the WZS pig genome. After trimming and quality control, yielding 155,075,433,268 of clean Illumina bases (∼60× coverage) (Supplementary Table S1), 141.49 Gb of HiFi data (∼53× coverage; average length ∼19 Kb), and 326.05 Gb of ONT data (N50 length ∼81 Kb; ∼125× coverage; average length ∼25 Kb), and 476.39 Gb of Hi-C data (∼183.23× coverage). The draft assembly spanned ∼2.78 Gb, with an N50 of 135,65 Mb, comprising 107 contigs within 79 unmatched contigs (∼103 Mb) (Supplementary Table S2). After eliminating redundancy, and anchoring with Hi-C data for gap filling and telomere repair, the final assembly yielded a ∼2.68 Gb genome size organized to 20 chromosomes (Fig. 1A and 1B), with an N50 length of ∼142.87 Mb (Supplementary Table S3 and S4). The sequencing reads were back aligned to WZS-T2T, achieving mapping rates of 99.96% for Illumina reads, 99.92% for HiFi data, and 96.98% for ONT data. QV of WZS-T2T was 70.2 value, higher than 51.41 of T2T-pig1.0. GCI values were 84.67 for both ONT data and HiFi data, comparable to the human genome CHM13 V2.0 (GCI = 87.04) (Supplementary Table S5).

**Figure 1:**
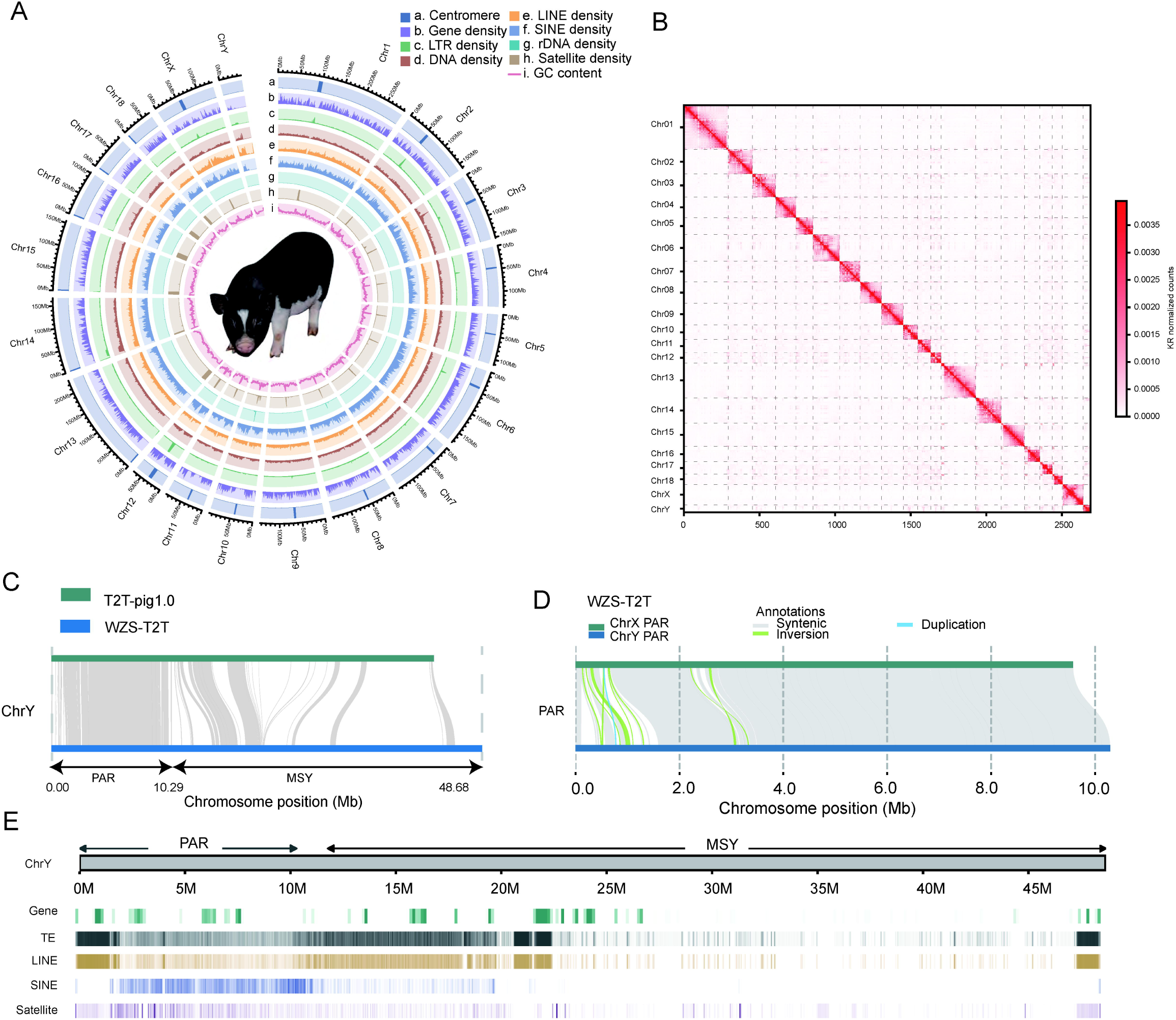
Genome description of WZS pig. (A) The Genomic landscape of WZS pig. Tracks a-i represented Centromere (a), Gene density (b), LTR density (c), DNA TRANSPOSONS density (d), LINE density (e), SINE density (f), rDNA TRANSPOSONS density (g), Satellite density (h), GC content (i). (B) Hi-C interaction map. X-axis presented the distance of chromosomes, and Y-axis presented the 20 chromosomes. (C) Syntenic regions on Y chromosomes between two WZS pig genomes (WZS-T2T, T2T-pig1.0). (D) The comparison of SVs in PAR between X and Y chromosomes. Top block was PAR in X chromosome, and the below block was PAR in Y chromosome of WZS-T2T. The color lines between two pigs presented different types of SVs (synteny, inversion, and duplication). PAR, pseudoautosomal region. SVs, structural variations. (E) Gene and repeat sequences (TE, LINE, SINE, satellite) across the PAR and MSY.

### Genome annotation

Repeat sequences of WZS-T2T comprised 43.74% of the genome, spanning a total length of ∼1.17 Gb across 3,708,253 sequences, among which long interspersed nuclear elements (LINEs) (23.25%) and short interspersed nuclear elements (SINEs) (10.94%) were the most prevalent (Supplementary Table S6-S8), while repeat sequences of T2T-pig1.0 accounted for 48.25% of the genome, spanning ∼1.27 Gb, with LINEs being the most abundant (24.46%). Furthermore, a total of 23,100 protein-coding genes was predicted using *ab initio* and homology-based approaches (Supplementary Table S9 and S10), and functional annotation identified 22,693 known genes based on the KEGG, NR, Uniprot, GO, KOG, Pfam, and InterPro databases (Supplementary Table S11). The BUSCO analysis indicated 99.4% completeness of WZS-T2T, which was higher than other available pig genomes, especially higher than 97.9% of T2T-pig1.0 (Supplementary Table S4).

### Telomeres and centromeres

Telomeric repeats of WZS-T2T were identified at the both ends of all the 20 chromosomes, ranging from 125 to 3,089 copies, and the most repeat copies of 3,089 was at the downstream of chromosome 17. The number of average repeat sequences was 996 copies at the upstream ends, and 1,095 copies at the downstream ends, while the repeat sequences > 1,000 copies on at least one end of 15 chromosomes. The Y chromosome carried 705 telomeric repeats upstream and 229 downstream, further confirming assembly completeness. (Supplementary Table S12). Similarly, T2T-pig1.0 assembled telomeres for 20 chromosomes. Candidate centromeric regions were identified by clustering tandem repeats, and satellite DNA 336 (336 bp) was the predominant repeat monomer among them. Centromeres were detected at the metacentric or submetacentric positions of all the 18 autosomes and the X chromosome (Supplementary Fig. S1 and Supplementary Table S13), with the acrocentric presented on chromosome 13-18, which was similar to T2T-pig1.0 [14].

### Y chromosome comparisons between pig breeds

The Y chromosome (ChrY) showed great diversity in length and genes of different pig breeds. Firstly, ChrY of four available T2T pig genomes were compared in this study. In detail, three pig breed genomes, including WZS-T2T, T2T-pig1.0, Jinhua pig, Chenghua pig, have assembled an ChrY with one contig, which measured ∼48.68 Mb, ∼43.24 Mb, ∼43.10 Mb and ∼29.24 Mb, respectively (Supplementary Table S14), indicating that WZS-T2T assembled the longest ChrY, which contributed to the comprehensive comparison of genetic characteristics of porcine ChrY. WZS-T2T, T2T-pig1.0, Jinhua pig, Chenghua pig harbored 79, 111, 188, and 47 protein-coding genes, respectively. The collinearity among the ChrY of the three pig breeds showed that 46 genes shared homology between WZS and Jinhua pigs (Fig. S2), seven genes between WZS and Chenghua pigs, and 12 genes between two WZS pigs, indicating abundant specific genes were existed in ChrY among different pig breeds. Moreover, the repeat sequences of ChrY from WZS-T2T comprised 94.08% (∼45.80 Mb) of ChrY, while T2T-pig1.0 accounted for 84.9% (∼36.72 Mb) of ChrY. Furthermore, both pseudoautosomal region (PAR) and male-specific region of the Y chromosome (MSY) were analyzed. PAR of WZS-T2T and T2T-pig1.0 presented strong synteny with each other, and MSY revealed less synteny (Fig. 1C and Supplementary Table S15), suggesting the diversity of ChrY even within the same breed. PAR of WZS-T2T spanned ∼9.58 Mb on the ChrX and ∼10.29 Mb on the ChrY, with strong sequence homology (Fig. 1D and Supplementary Table S16), which was similar to ∼10.2 Mb of PAR on the ChrY of T2T-pig1.0. The repeat sequences of PAR of ChrX and PAR of ChrY from WZS-T2T accounted for 16.59% (∼8.08 Mb) and 18.56% (∼9.04 Mb) of the ChrY, respectively (Supplementary Table S17). The male-specific region of the Y chromosome (MSY) of WZS-T2T spanned ∼36.76 Mb, with ∼75.52% repeat sequences of ChrY (Fig. 1E and Supplementary Table S18), and LINEs (∼8.41 Mb) was dominant. A number of 42 genes and 37 genes were on the PAR of ChrX and PAR of ChrY in WZS-T2T, respectively, while 62 genes in T2T-pig1.0. In detail, PAR of ChrX and PAR of ChrY from WZS-T2T included nine homologous genes (Supplementary Table S19). Among these, *PLCXD1* (phosphatidylinositol-specific phospholipase C X domain containing 1) [71] and *SHOX* (SHOX homeobox) [72] are involved in spermatogenesis. MSY of WZS-T2T harbored 33 genes (Supplementary Table S20), while T2T-pig1.0 included 49 genes. MSY mainly harbored spermatogenesis-related genes of *SRY* (sex determining region Y) [73], *USP9Y* (ubiquitin specific peptidase 9 Y-linked), *UTY* (ubiquitously transcribed tetratricopeptide repeat containing, Y-linked). Furthermore, eleven genes were shared by ChrY between WZS-T2T and T2T-pig1.0 (Supplementary Table S21), among which eight genes were located in PAR regions, and three genes in MSY regions. Among these eleven genes, *SHOX*, *PRKX* (protein kinase cAMP-dependent X-linked catalytic subunit), and *DDX3X* (DEAD-box helicase 3 X-linked), located in the PAR regions, as well as *SRY* in the MSY region, are associated with sex differentiation, whereas the energy metabolism gene *SLC25A4* (solute carrier family 25 member 4) and the macrophage-related receptor gene *CSF2RA* (colony stimulating factor 2 receptor subunit alpha) were specific to the PAR regions on ChrY of WZS-T2T. Therefore, the porcine ChrY demonstrates significant inter-individual variation in both length and gene composition even within the same breed, indicating that fully resolving its genetic landscape will require complete assemblies of ChrY from a diverse set of individuals.

### SVs comparisons between WZS-T2T and T2T-pig1.0

Relative to T2T-pig1.0, WZS-T2T included 10,868,902 SNPs, and 26,269 SVs (>50 bp) (Supplementary Table S22). The SVs were further classified into 705 transitions, 200 inversions, 170 duplications, 12,687 insertions, 12,507 deletions (Fig 2A and Supplementary Table S23). Functional annotation revealed that 3,049,576 of these SNPs were located in genes, 73,484 in exonic regions (Supplementary Table S24), and 3,056 of these SVs were located in exonic regions, 631,735 in intronic regions and 1,572,038 in intergenic regions (Supplementary Table S25). GO and KEGG enrichment analyses of the SV-affected genes showed significant enrichment in cytoskeleton pathways and MAPK signaling pathway (Fig. 2B). Notably, an inversion SV on chromosome 10 (Chr10) with range 0 bp to 33.86 Mb was identified between WZS-T2T and T2T-pig1.0 (Fig. S3). The sequence orientation of the Chr10 of WZS-T2T was verified by three proofs. First, IGV snapshots of ONT and HiFi reads demonstrated the reads spanned the joint sites of SV. Second, collinearity analysis showed that sequence orientations of SV on Chr10 were similar among five assemblies of WZS-T2T, Jinhua pigs, Sscrofa 11.1, Min and Rongchang pigs, whereas T2T-pig1.0 displayed reversed orientation with the five assemblies (Fig. 2C). Furthermore, T2T-pig1.0 also displayed SVs of varying lengths with other 14 pig assemblies (Bannamini pig, Ningxiang pig, Tunchang pig, Shawutou pig, Korean mini pig, Babraham pig, Landrace pig, US corssbred pig, Chenghua pig, Tongcheng pig, Meishan pig_Liu, Laiwu pig, Bamaxiang pig, Huai pig) [14], but aligned to only four assemblies (Bamei pig, Juema pig, Diannanxiaoer pig, Meishan_Li); therefore, an inversion SV on Chr10 were identified between T2T-pig1.0 and 19 assemblies, which suggested that the sequence orientation of SV on Chr10 of T2T-pig1.0 showed prominent differences with ∼83% (19/23) of the available pig assemblies, in contrast, sequence orientation of SV on Chr10 of WZS-T2T presented highly consistent with ∼83% of the available assemblies. Third, telomeres are expected to be positioned at the ends of chromosomes, and the telomeres position (2 bp to 19,048 bp) involved in the inversion SV was at one end of Chr10 for of WZS-T2T (Fig. 2C). Consequently, these three proofs identified the accuracy of sequence orientation for SV on Chr10 of WZS-T2T. Furthermore, this inversion SV on Chr10 included 221 and 241 genes for WZS-T2T and T2T-pig1.0, respectively, and shared 85 genes with opposite chromosomal orientations in two WZS pigs (Supplementary Table S26), which likely reflects prominent porcine individual-level variation and the necessity of obtaining assemblies from a broader range of WZS pig individuals.

**Figure 2:**
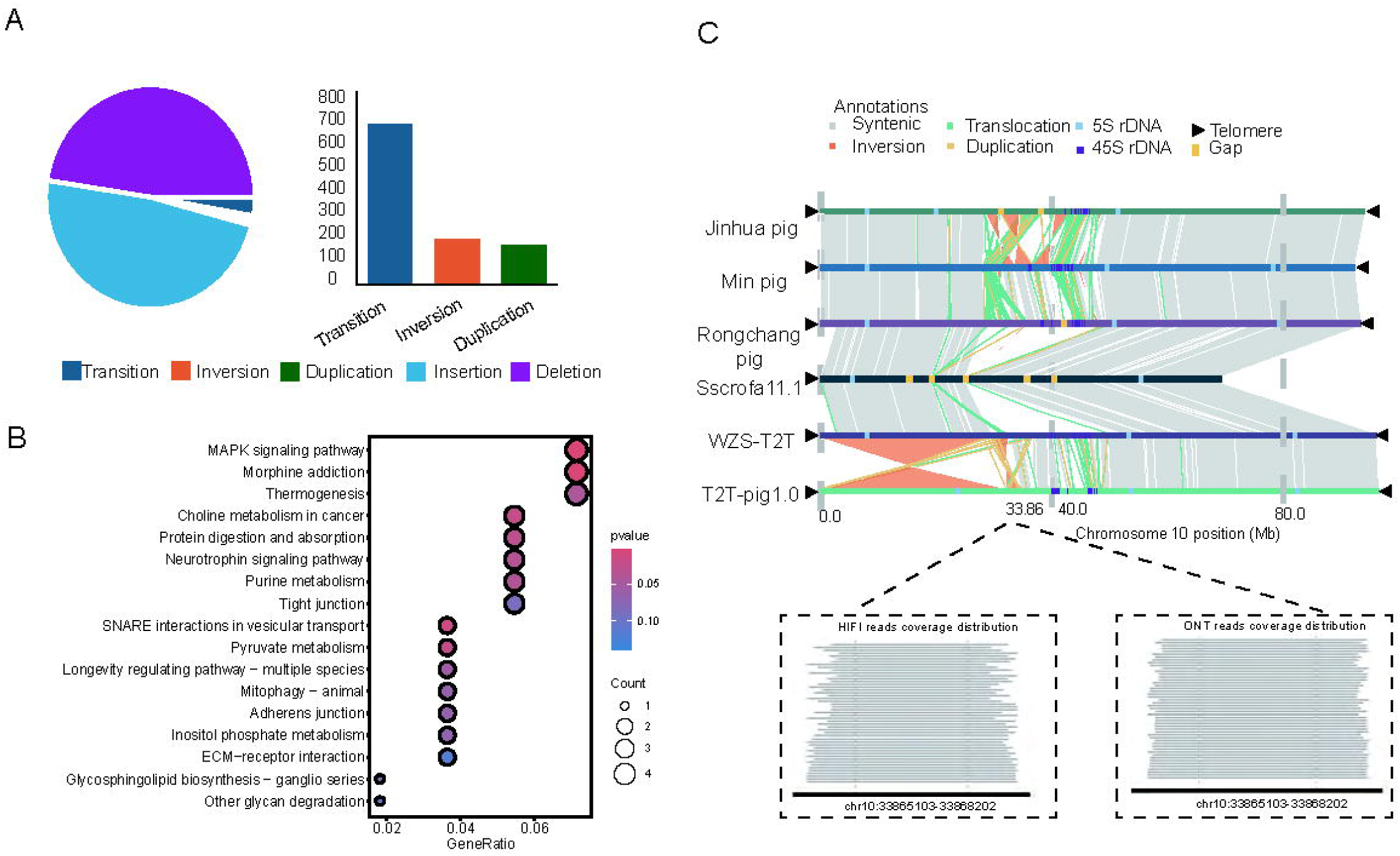
The comparisons of SVs from two WZS pigs. (A) The types and numbers of SVs. (B) The KEGG function enrichment of SVs. X-axis presented the gene number of each KEGG pathway, and Y-axis presented the name of KEGG pathway. (C) Chr10 synteny analysis and reads coverage. The pig breeds from top to bottom were Jinhua pig, Min pig, Rongchang pig, Sscrofa 11.1, WZS-T2T, T2T-pig1.0. The collinearity blocks on Chr10 were consistent among WZS-T2T, Min and Rongchang pigs, Jinhua pigs, and Sscrofa 11.1, but not with the T2T-pig1.0. The black triangles at the ends of chromosomes are telomeres. The HiFi and ONT reads both spanned the SV joint point at ∼33.86 Mb.

### Expansion and contraction of gene families

Orthologous groups (OGs) across 196,065 genes from 6 species representing two pig breeds were counted, and phylogenetic tree was constructed. As expected, the WZS and Duroc pig breeds were closely related, and diverged approximately at 3.3 million years ago (MYA) (Fig. 3A), which fell within 95% confidence interval of the divergence time (0.8 MYA - 5.5 MYA) between Duroc and T2T-pig1.0 [14]. In total, 184,568 genes were clustered into 20,361 orthologous groups (OGs), leaving 11,497 unassigned. Of these, 5,431 single-copy OGs were shared by among all species and breeds (Fig. 3B). WZS-T2T presented 558 and 1,315 gene families for expansion and contraction (Supplementary Table S27 and S28), while T2T-pig1.0 expanded and contracted for 541 and 420 gene families, respectively. The expansion gene families were involved in positive regulation of immature T cell proliferation and interleukin-1 receptor binding (Fig. 3C), while the contraction gene families mainly regulated carbohydrate metabolic process, and carbohydrate metabolic process (Fig. 3D). Additionally, 2,054 positively selected genes (PSGs) were identified, contributing to adaptation in response to environmental fluctuations. These PSGs were primarily involved in positive regulation of calcium-mediated signaling, Wnt signaling pathway, PI3K-Akt signaling pathway, and melanogenesis.

**Figure 3:**
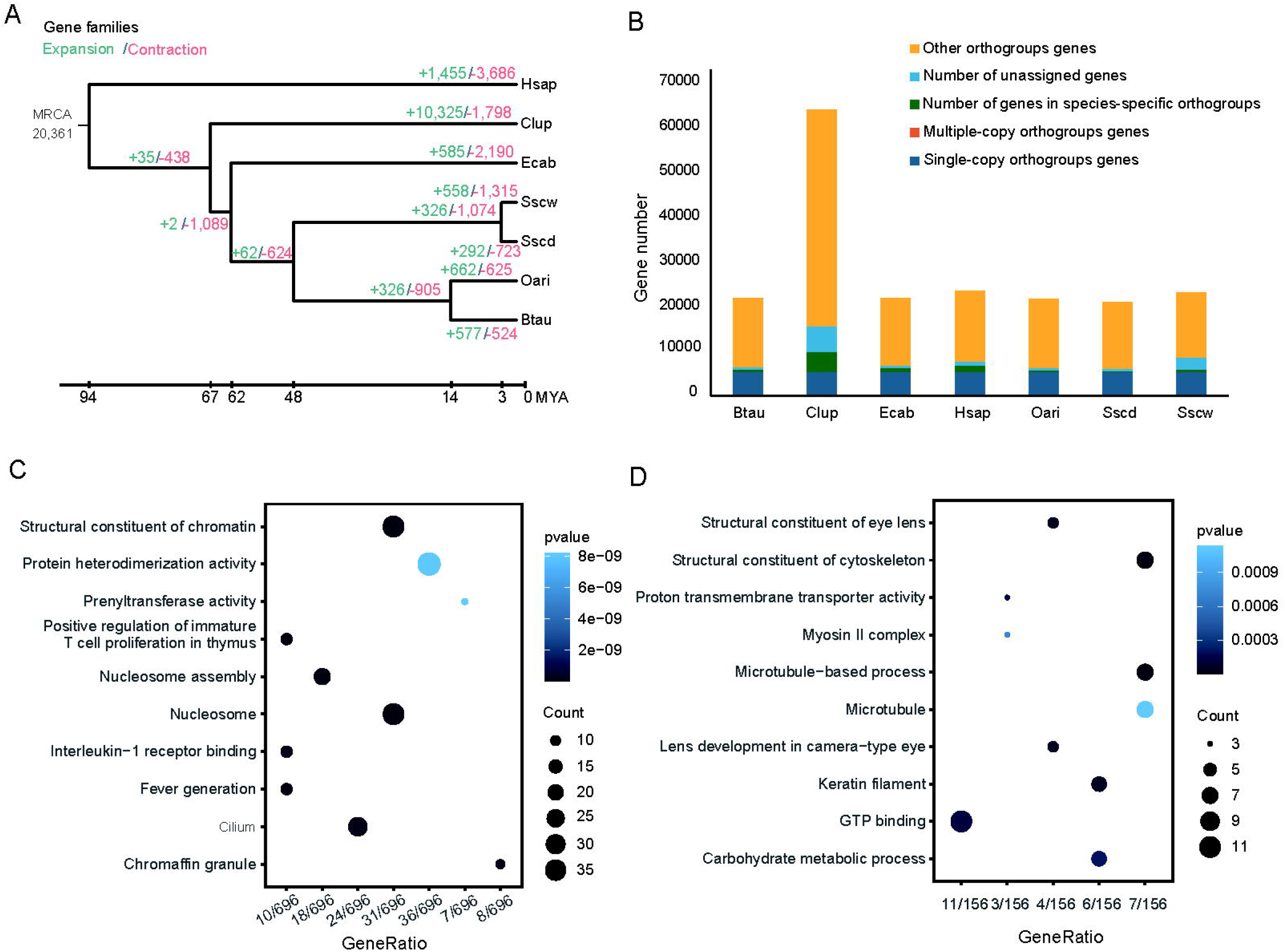
The expansion and contraction of gene families. (A) Phylogenetic tree of 6 species, including 2 pig breeds, constructed using single-copy orthologous genes. Numbers of contracted and expanded gene families are marked on divergent nodes. Green color words presented expansion genes number, and red color words presented contraction genes number. The 6 species were: Oari: Ovis aries; Btau: Bos taurus; Sscw: *Sus scrofa* WZS pig; Sscd: *Sus scrofa* Duroc Sscrofa 11.1; Ecab: Equus caballus; Hsap: Homo sapiens; Clup: Canis lupus. (B) The gene numbers in different types of OGs. Btau, Bos taurus; Clup, Canis lupus; Ecab, Equus caballus; Hsap, Homo sapiens; Oari, Ovis aries; Sscw, *Sus scrofa* Wuzhishan pig; Sscd, *Sus scrofa* Duroc Sscrofa 11. OGs, orthologous groups. (C) The GO terms of genes in expansion gene families. X-axis presented the gene number ratio of GO terms, and Y-axis presented the name of GO terms. GO, Gene Ontology. (D) The GO terms of genes in contraction gene families.

### Population difference analysis

A total of ∼4,223.28 Gb sequencing data of 147 samples and ∼1,875.53 Gb download whole genome sequencing data (PRJNA994680) were used in this study (Supplementary Table S29). The pig breeds and the number of SNPs per population were: WZS pigs (19,957,874), Yuxi black pigs (YX pigs, 18,805,847), Tunchang pigs (TC pigs, 15,984,774), Large White pigs (LW, 12,407,372), and Duroc pigs (11,241,323). The majority of SNP density near the ends of chromosomes is greater than that in the middle of the chromosomes (Supplementary Fig. S4), and the SNPs were the largest in the intergenic regions (Supplementary Table S30). Among these five pig populations examined in this study, WZS pigs exhibited the highest genetic diversity (π = 2.699E-03), while Duroc pigs was the lowest (π = 1.624E-03) (Supplementary Table S31). The lowest pairwise *F_ST_* value was observed between WZS-pigs and TC pigs (*F_ST_* = 0.103), while the highest was between TC and Duroc, and between TC and LW pigs (*F_ST_* = 0.329) (Supplementary Table S32). The phylogenetic tree separated the five pig breeds (Fig. 4A), and the PCA revealed similar population groups (Fig. 4B). These results indicate consistent population classifications. Population structure analysis divided the samples into five groups when K = 5 (Fig. 4C). The tendency of LD was consistent with the π value. Linkage disequilibrium (LD) decay analysis showed that Duroc pigs had the highest LD levels, reflecting strong domestication and selective pressure, while WZS pigs exhibited the lowest LD (Fig. 4D), suggesting WZS pigs have a shorter domestication history and a slower LD decay compared to other four populations, which was consistent with genetic diversity patterns. As a whole, the genome research was crucial for uncovering both inter-breed genetic differences and substantial variation even within the same breed, and revealing potential advantages of both agricultural production and biomedical targets for WZS pigs.

**Figure 4:**
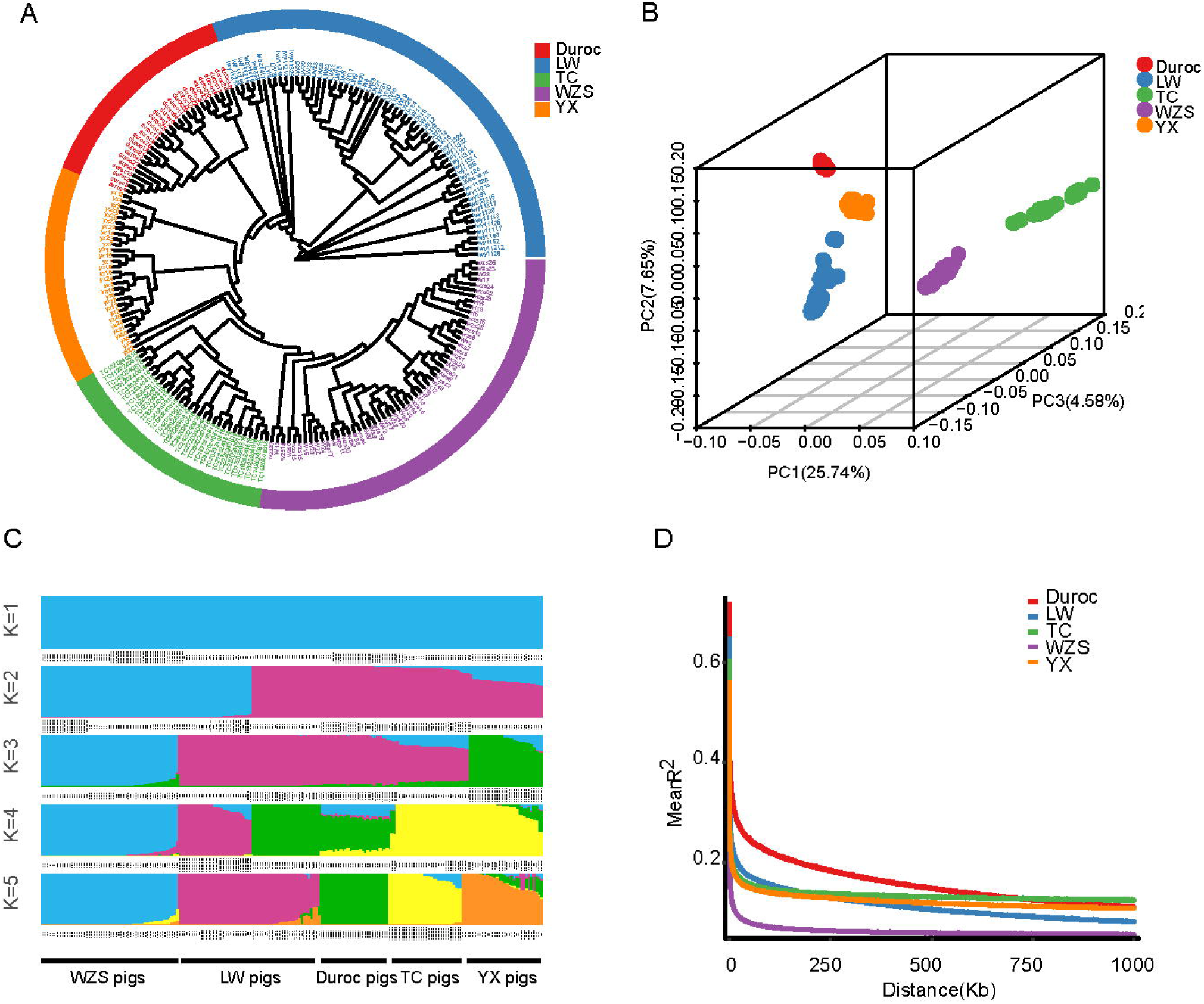
Population differentiation analysis. (A) The phylogenetic tree of five pig breeds. LW, Large White pigs; TC, Tunchang pigs; WZS, Wuzhishan pigs; YX, Yuxi black pigs. (B) The PCA of five pig breeds. (C) Admixture analysis of five populations. (D) The LD value of five pig breeds. X-axis presented the physics distances. Y-axis presented the LD coefficient R^2^.

## Discussion

Both pig genome assemblies (WZS-T2T and T2T-pig1.0) achieved high quality, while this study revealed the similarities and differences between them. The genome size and N50 length were similar between two WZS pig assemblies, but the genome quality of WZS-T2T was higher. WZS-T2T assembly yielded a ∼2.68 Gb genome size organized to 20 chromosomes, with an N50 length of ∼142.87 Mb, which was similar to that of T2T-pig1.0 (genome size ∼2.6 GB, N50 ∼144.9 Mb) [14]. However, QV (70.2) of WZS-T2T was higher than 51.41 of T2T-pig1.0. For comparison of annotation results, repeat sequences were accounted for 43.74% and 48.25% of WZS-T2T and T2T-pig1.0 genomes, respectively, and LINEs were the most prevalent for both WZS pigs. The protein-coding genes number 23,100 of WZS-T2T was a little more than 21,713 of T2T-pig1.0, and was similar to that of Jinhua pig (23,924) [11], and Chenghua pig (23,453) [12]. BUSCO value of WZS-T2T (99.4%) were also higher than 97.9% of T2T-pig1.0. Telomeres of two WZS pig genomes were all assembled on 20 chromosomes, and the acrocentric presented on chromosome 13-18 for both WZS pigs.

The diversity of ChrY were compared between two WZS pigs. The length of ChrY of WZS-T2T (∼48.68 Mb) was the longest among the available pig breeds assembled with Y chromosomes. PAR of WZS-T2T and T2T-pig1.0 presented strong synteny with each other, and MSY revealed more diversity regions, and the length of PAR on ChrY of WZS-T2T (∼10.29 Mb) was similar to that on ChrY of T2T-pig1.0 (∼10.2 Mb). The collinearity analysis showed that 12 genes of ChrY were shared by two pigs, and eight genes in PAR region, while three genes in MSY region. Most shared genes of ChrY between two WZS pigs were spermatogenesis-related genes, including *SHOX*, *PRKX*, and *DDX3X*, whereas energy metabolism gene *SLC25A4* and the macrophage-related receptor gene *CSF2RA* were specific to WZS-T2T, suggesting the nutrition metabolism and immune responses were associated with individual variation.

An inversion SV on Chr10 with range 0 bp to 33.86 Mb was identified between WZS-T2T and T2T-pig1.0, and three proofs were proposed to identify the accuracy sequence orientations of the inversion SV on Chr10 in WZS-T2T in this study. Notably, the T2T-pig1.0 also existed an inversion SV on Chr10 with Sscrofa 11.1, and two methods were used to identify the sequence orientation of this SV on T2T-pig1.0 by the author [14]. The two out of the three proofs used in this study were compared with two proofs in the published T2T-pig1.0 [14]. In detail, the first proof was that ONT and HiFi reads spanned the joint sites (33.86 Mb) of SV on Chr10 in WZS-T2T, which was similar to the method used to identify the inversion SV on Chr10 between T2T-pig1.0 and Sscrofa 11.1 [14]. The second proof was the same sequence orientation of SV among five assemblies of WZS-T2T and other four pig genomes, whereas the SV orientation in T2T-pig1.0 was opposite to that observed in the five assemblies. Furthermore, the collinearity analysis revealed an inversion SV on pig Chr10 between T2T-pig1.0 and other 14 pig assemblies, and this SV of T2T-pig1.0 aligned to only four assemblies [14], suggesting the orientation of this SV in T2T-pig1.0 was found to be reversed to a total of 19 pig assemblies with varying lengths, including five assemblies (WZS-T2T, Jinhua pig, Min pig, Rongchang pig, Sscrofa 11.1) from collinearity analysis of this study. As a result, the sequence orientation of this SV on Chr10 in the T2T-pig1.0 assembly is prominently different from that in ∼83% (19/23) of the available assemblies. Conversely, sequence orientation of SV on Chr10 of WZS-T2T presented highly consistent with ∼83% of the available assemblies. The third proof was that telomeres are expected to located at the ends of chromosomes, and the telomeres position (2 bp to 19,048 bp) involved in the inversion SV on Chr10 of WZS-T2T was at one end of Chr10. Therefore, three proofs, including the spanning joint sites by ONT and HiFi reads, high ratio (∼83%, 19/23) of similarity sequence orientation of SV on Chr10 among WZS-T2T and 23 pig assemblies, and the right position of telomeres on Chr10, identified the accuracy of sequence orientation of SV on Chr10 in WZS-T2T. However, the inversion SV on Chr10 between T2T-pig1.0 and 19 other assemblies was explained as a true SV rather than an assembly error based on the two proofs [14], the first was long reads spanning across the joint sites of SV, and the second was that SV sequence orientation of T2T-pig1.0 was consistent with only four out of the 23 pig assemblies (∼17%). The prominent differences between two WZS pigs indicated that the underlying causes of the inversion SV on Chr10 remain to be further in-depth explored.

Furthermore, the population difference analysis indicated that WZS pigs exhibited higher genetic diversity, and showed slower LD decay speed and shorter domestication time compared to other four populations (TC pigs, YX pigs, LW pigs and Duroc pigs) examined in this study. In detail, WZS pigs exhibited the highest π value, and the lowest pairwise *F_ST_* value was detected between WZS pigs and TC pigs, while the tendency of LD were consistent with the π value.

## Conclusions

The comparison results between two WZS pig genomes (WZS-T2T and T2T-pig1.0), including higher QV value, higher BUSCO value, longer ChrY length, and the reasonable sequence orientation of inversion SV on Chr10 for WZS-T2T, indicated WZS-T2T provided a higher-quality genome. Genes comparisons in ChrY showed that most of the spermatogenesis-related genes were shared by two WZS pigs, whereas energy metabolism gene *SLC25A4* and the macrophage-related receptor gene *CSF2RA* of ChrY were specific to WZS-T2T. Specifically, the less synteny of MSY region of ChrY revealed the huge individual-level variations. Although the accuracy sequence orientation of the inversion SV on Chr10 ranging from 0 bp to 33.86 Mb of WZS-T2T has been identified by three proofs, the underlying cause for the inversion SV is required in-depth investigation in the future. As a result, the genome assembly and population analysis were crucial for uncovering both inter-breed genetic differences and substantial variation even within the same breed, and revealing the great potential for agricultural production and medical models for WZS pigs.

## Additional Files

**Supplementary Figure S1.** Candidate centromere regions of WZS-T2T.

**Supplementary Figure S2.** The collinearity between the Y chromosomes of WZS-T2T and Jinhua pig.

**Supplementary Figure S3.** The syntenic analysis between WZS-T2T and T2T-pig1.0.

**Supplementary Figure S4.** The SNPs density of WZS pigs.

**Supplementary Table S1.** Statistics of sequencing data of WZS-T2T.

**Supplementary Table S2.** The initial assembly with gaps of WZS-T2T

**Supplementary Table S3.** The statistics of chromosomes of WZS-T2T

**Supplementary Table S4.** Assessment of different pig breeds genome sequences.

**Supplementary Table S5.** The GCI of 20 chromosomes of WZS-T2T.

**Supplementary Table S6.** The annotation of transposable elements sequences of WZS-T2T.

**Supplementary Table S7.** The annotation of tandem repeat sequences of WZS-T2T.

**Supplementary Table S8.** The annotation of non-coding RNA sequences of WZS-T2T

**Supplementary Table S9.** The annotation of gene structures of WZS-T2T.

**Supplementary Table S10.** The length of protein-coding genes of WZS-T2T.

**Supplementary Table S11.** The comparison of annotation of protein-coding genes of pigs.

**Supplementary Table S12.** The repeat sequences of telometers of WZS-T2T.

**Supplementary Table S13.** The prediction of centromeres of WZS-T2T.

**Supplementary Table S14.** The length of chromosome Y in six pigs genome.

**Supplementary Table S15.** The synteny of Y chromosoe between WZS-T2T and T2T-pig1.0.

**Supplementary Table S16.** The comparison of PAR between X and Y chromosomes in WZS-T2T.

**Supplementary Table S17.** The sequence types of PAR of X and Y chromosomes of WZS-T2T.

**Supplementary Table S18.** The identification of MSY region on the Y chromosome of WZS-T2T.

**Supplementary Table S19.** Homologous genes of ChrX PAR and ChrY PAR of WZS-T2T.

**Supplementary Table S20.** The repeat sequences of MSY region on the Y chromosome of WZS-T2T.

**Supplementary Table S21.** Shared genes in ChrY of WZS-T2T and T2T-pig1.0.

**Supplementary Table S22.**Variations comparison between WZS-T2T and T2T-pig1.0.

**Supplementary Table S23.** The SVs types of WZS-T2T and T2T-pig1.0.

**Supplementary Table S24.** The structure annotation of SNPs of WZS-T2T and T2T-pig1.0.

**Supplementary Table S25.** The structure annotation of SVs of WZS-T2T and T2T-pig1.0.

**Supplementary Table S26.** The genes on Chr10 shared by both WZS-T2T and T2T-pig1.0.

**Supplementary Table S27.** The genes in expansion gene families of WZS-T2T.

**Supplementary Table S28.** The genes in contraction gene families of WZS-T2T.

**Supplementary Table S29.** The summary of whole genome sequencing data from five populations.

**Supplementary Table S30.** The distribution of SNPs of five populations.

**Supplementary Table S31.** The genetic diversity of five populations.

**Supplementary Table S32.** The pairwise FST between WZS pigs and other pig breeds.

## Abbreviations

T2T: telomere-to-telomere
WZS: Wuzhishan
WZS-T2T: WZS pig genome assembled in this study
ChrY: Y chromosome
SHOX: SHOX homeobox
PRKX: protein kinase cAMP-dependent X-linked catalytic subunit
DDX3X: DEAD-box helicase 3 X-linked
SRY: sex determining region Y
SLC25A4: solute carrier family 25 member 4
CSF2RA: colony stimulating factor 2 receptor subunit alpha
Chr10: chromosome 10
GGTA1: glycoprotein alpha-galactosyltransferase 1
SVs: structural variations
HiFi: High-Fidelity Sequencing
ONT: Oxford Nanopore Technologies
Ccs: circular consensus sequencing
Hi-C: high-throughput/resolution chromosome conformation capture
GCI: Genome Continuity Inspector
BUSCO: Benchmarking Universal Single-Copy Orthologs
TEs: transposable elements
TRs: tandem repeats
LTRs: long terminal repeats
TIRs: terminal inverted repeats
SSRs: simple sequence repeats
LDM: longissimus dorsi muscle
NR database: Non-Redundant Protein Sequence Database
GO: Gene Ontology
KEGG: Kyoto Encyclopedia of Genes and Genomes
OG: orthologous genes
ML: maximum likelihood
PARs: pseudoautosomal regions
SNPs: single nucleotide polymorphisms
MSY: male-specific regions on Y
TC: Tunchang pig
YX: Yuxi black pig
LW: Large White pig
IBS: identity-by-state
FST: Weir and Cockerham’s fixation index
LD: linkage disequilibrium
LINEs: long interspersed nuclear elements
SINEs: short interspersed nuclear elements
PLCXD1: phosphatidylinositol-specific phospholipase C X domain containing 1
USP9Y: ubiquitin specific peptidase 9 Y-linked
UTY: ubiquitously transcribed tetratricopeptide repeat containing, Y-linked
MYA: million years ago
PSGs: positively selected genes.

## Data availability

The WZS pig genome assembly have been released in the Genome Warehouse at the National Genomics Data Center, Beijing Institute of Genomics, Chinese Academy of Sciences/China National Center for Bioinformation, under accession number GWHHNIC00000000.1. The raw sequencing data for genome assembly and whole genome sequencing data have been deposited in the Genome Warehouse at the National Genomics Data Center, under accession number PRJCA036066 andPRJCA054653. The annotation files were stored in figshare at https://figshare.com/s/57033524d6fcec0afdc6.

## Declarations

The study was performed according to the Regulations for the Administration of Affairs Concerning Experimental Animals (Ministry of Science and Technology, China, revised on 1 March 2017), and complied with the Institutional Animal Care and Use Committee at the Experimental Animal Center of Hainan Academy of Agricultural Science (HNXMSY-20210503, revised on 3 May 2021).

## Author Contributions

Yuwei Ren, Zhe Chao, Xinjian Li and Guangliang Liu: conceived and supervised the study. Yuwei Ren: performed data analysis, wrote the original manuscript. Yuwei Ren, Zhe Chao, Feng Wang, Ruiping Sun, Xinli Zheng, Ruiyi Lin, Xuyang Lu, Yan Zhang, Linlin Chen, Wenshui Xin, Yanli Fei: collected samples and extracted DNA sampling. Yuwei Ren and Zhe Chao: revised the manuscript. All authors read and approved the final manuscript.

