## Supplemental Figures and Tables for "A telomere-to-telomere (T2T) pig genome assembly reveals Y chromosome diversity and structural variations of Wuzhishan pigs": Supplementary_Material.docx

**Supplementary Figures**

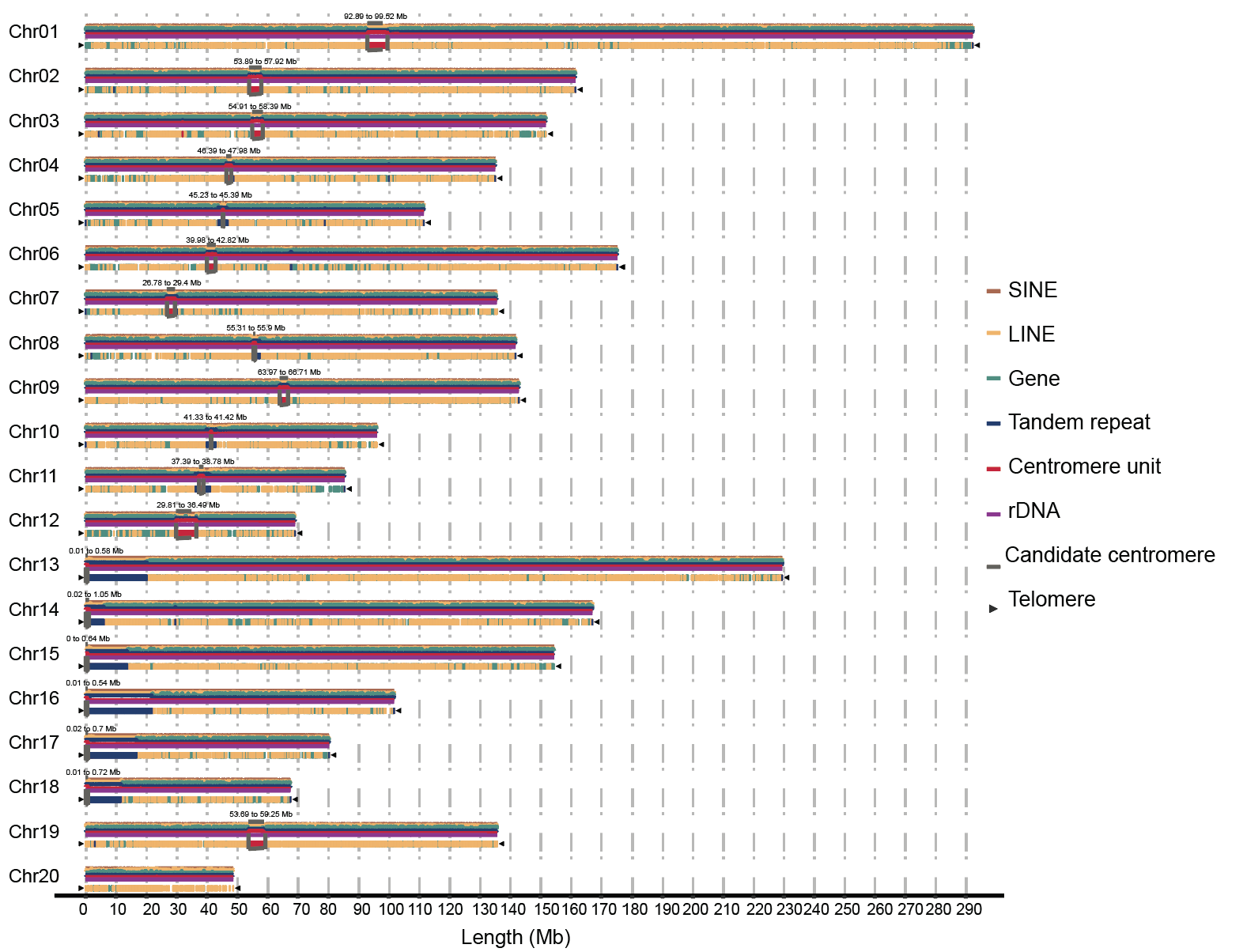

**Fig. S1** **Candidate centromere regions of WZS-T2T**

The candidate centromeres were marked as numbers at corresponding positions.

Chr1: 92.89 to 99.52 Mb, Chr2: 53.89 to 57.92 Mb, Chr3: 54.91 to 58.39 Mb, Chr4: 46.39 to 47.98 Mb, Chr5: 45.23 to 45.39 Mb, Chr6: 39.98 to 42.82 Mb, Chr7: 26.78 to 29.4 Mb, Chr8: 55.31 to 55.9 Mb, Chr9: 63.97 to 66.71 Mb, Chr10: 41.33 to 41.42 Mb, Chr11: 37.39 to 38.78 Mb, Chr12: 29.81 to 36.49 Mb, Chr13: 0.01 to 0.58 Mb, Chr14: 0.02 to 1.05 Mb, Chr15: 0 to 0.64 Mb, Chr16: 0.01 to 0.54 Mb, Chr17: 0.02 to 0.7 Mb, Chr18: 0.01 to 0.72 Mb, ChrX: 53.69 to 59.25 Mb.

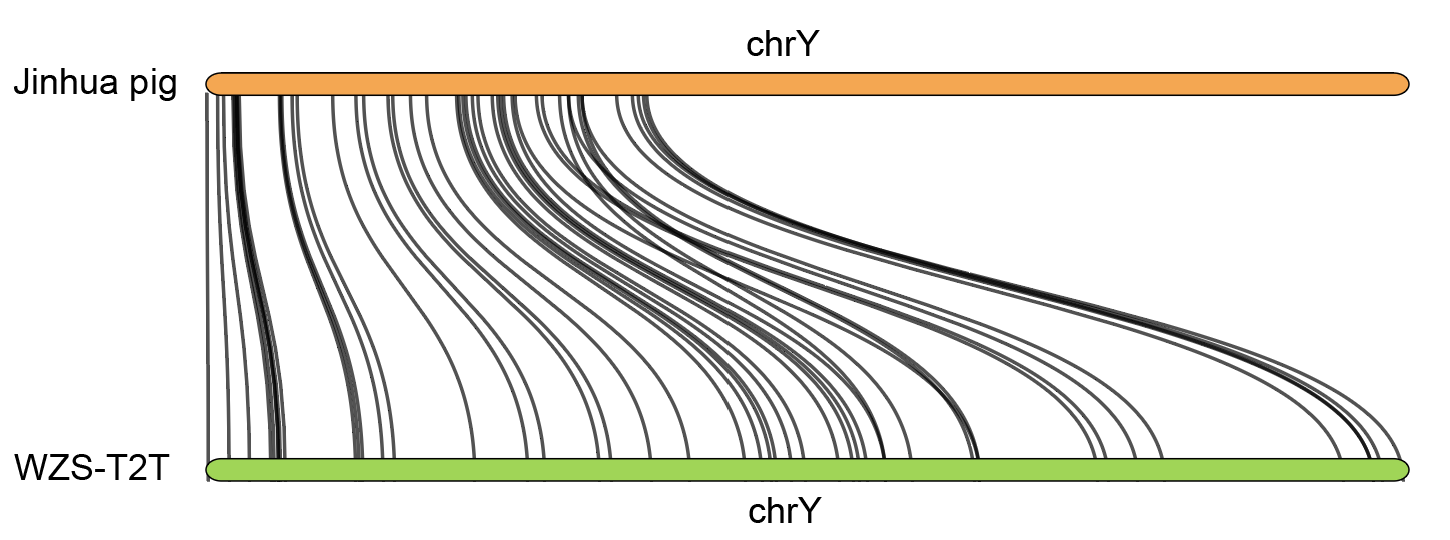

**Fig. S2** **The collinearity between the Y chromosomes of WZS-T2T and Jinhua pig**

The line between two pig breeds presented the collinearity of Y chromosome.

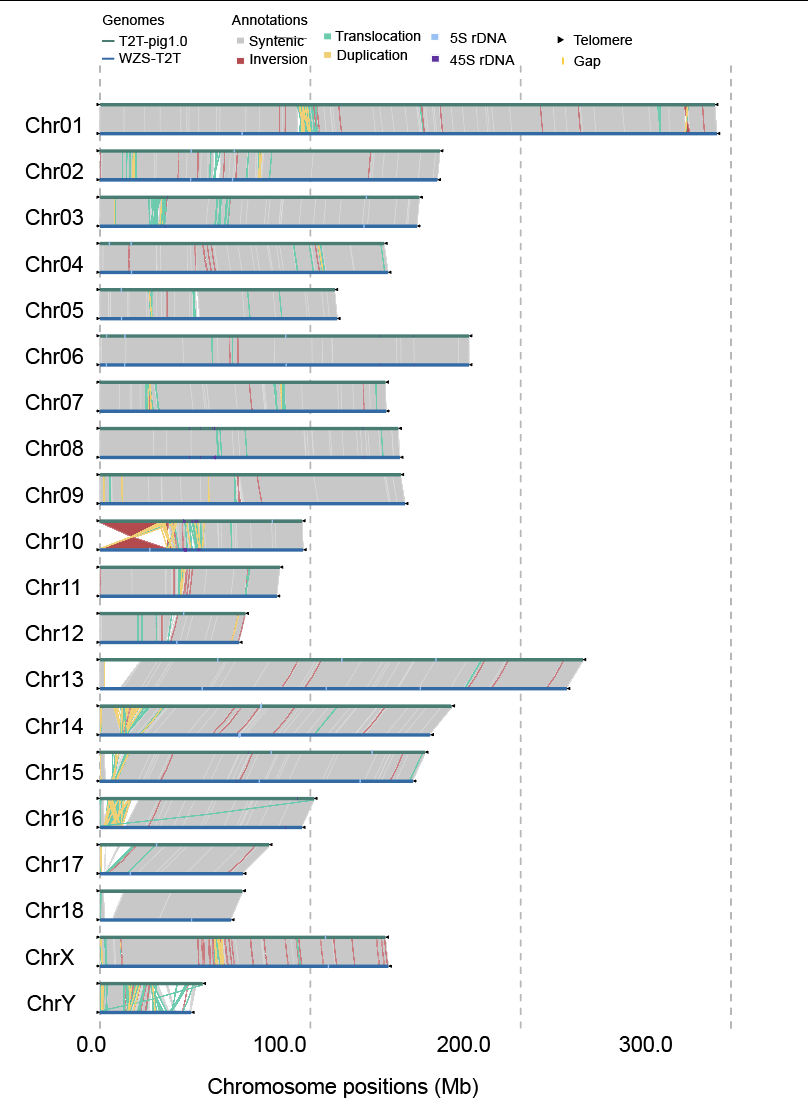

**Fig. S3** **The syntenic analysis between WZS-T2T and T2T-pig1.0**

The upper block was WZS-T2T, and the below block was T2T-pig1.0. The black triangles at the ends of chromosomes are telomeres, and the line between two pigs were the different types of SVs. SVs, structural variants.

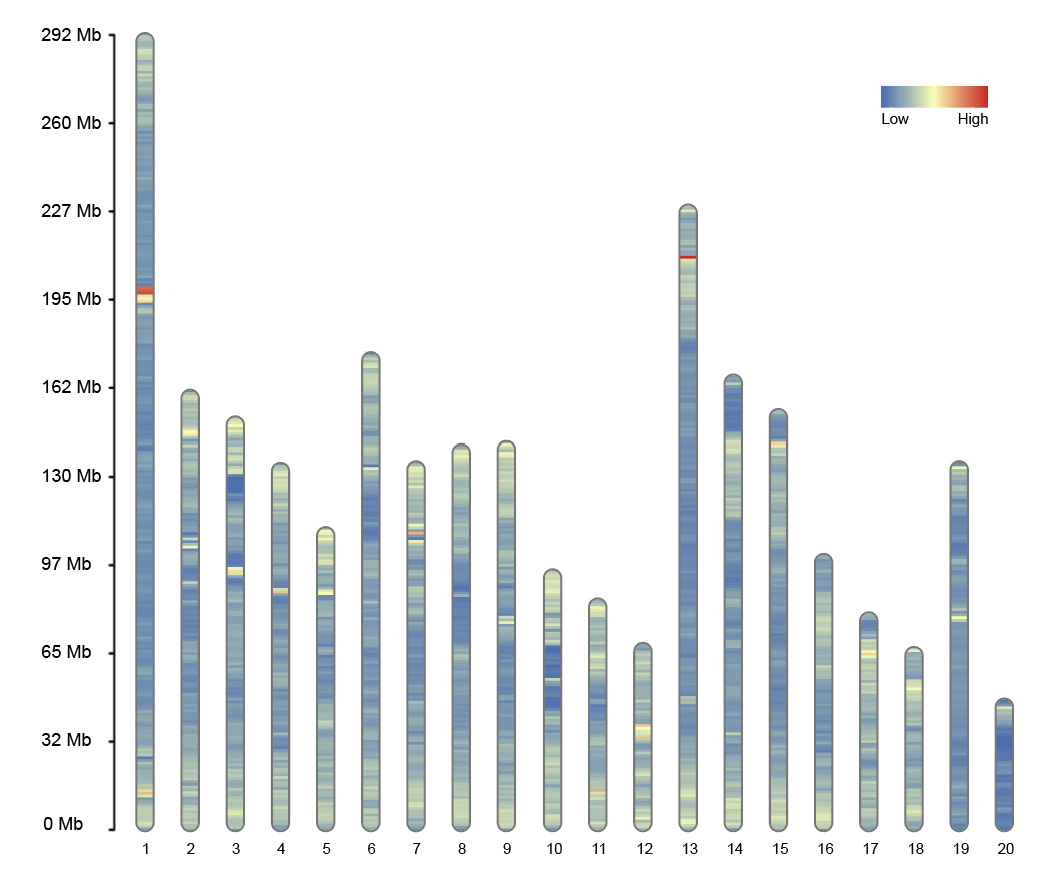

**Fig. S4 The SNPs density of WZS pigs**

The number below the column presented the 20 chromosomes, and the number on the left scale presented the length of chromosomes.
